# An osmolality/ salinity-responsive enhancer 1 (OSRE1) in intron 1 promotes salinity induction of tilapia glutamine synthetase

**DOI:** 10.1101/2020.03.16.994392

**Authors:** Chanhee Kim, Dietmar Kültz

**Affiliations:** Biochemical Evolution Laboratory, Department of Animal Sciences, University of California, Davis, CA, 95616

## Abstract

Euryhaline tilapia (*Oreochromis mossambicus*) are fish that tolerate a wide salinity range from fresh water to >3x seawater. Even though the physiological effector mechanisms of osmoregulation that maintain plasma homeostasis in fresh water and seawater fish are well known, the corresponding molecular mechanisms that control switching between hyper- (fresh water) and hypo-osmoregulation (seawater) remain mostly elusive. In this study we show that hyperosmotic induction of *glutamine synthetase* represents a prominent part of this switch. Proteomics analysis of the *O. mossambicus* OmB cell line revealed that glutamine synthetase is transcriptionally regulated by hyperosmolality. Therefore, the 5’ regulatory sequence of *O. mossambicus glutamine synthetase* was investigated. Using an enhancer trapping assay, we discovered a novel osmosensitive mechanism by which intron 1 positively mediates *glutamine synthetase* transcription. Intron 1 includes a single, functional copy of an osmoresponsive element, osmolality/salinity-responsive enhancer 1 (OSRE1). Unlike for conventional enhancers, the hyperosmotic induction of *glutamine synthetase* by intron 1 is position dependent. But irrespective of intron 1 position, OSRE1 deletion from intron 1 abolishes hyperosmotic enhancer activity. These findings indicate that proper intron 1 positioning and the presence of an OSRE1 in intron 1 are required for precise enhancement of hyperosmotic *glutamine synthetase* expression.

## Introduction

Euryhaline fish have evolved the capacity to utilize a suite of osmoresponsive genes for rapidly switching between hypo- and hyper-osmoregulation in response to salinity stress to maintain plasma ionic and osmotic homeostasis^1^. Mozambique tilapia (*O. mossambicus*) are representative euryhaline fish belonging to the family of cichlidae, which consists of many species that are uniquely adapted to specific environments^2,3^. A remarkable adaptive trait of *O. mossambicus* is its ability to tolerate large and rapid salinity fluctuations^4^. This species has evolved molecular mechanisms for rapidly turning on and off genes that encode enzymes and transporters involved in hypo- and hyper-osmoregulation^5–7^. However, the regulatory and evolutionary mechanisms controlling environmental (e.g. salinity, temperature, and hypoxia) regulation of gene expression in fish are still largely elusive. This lack of knowledge contrasts with the evolutionary diversity of fish, which have radiated into virtually any aquatic ecological niche. Previous studies investigating which parts of the genome have a functional role in the evolution of organisms have stressed *cis*-regulatory elements (CREs) as major targets of evolutionary adaptation^8,9^. Therefore, alterations of CREs are considered potent drivers of evolutionary adaptation^10^.

CREs typically contain binding sites for transcriptional regulators that orchestrate gene expression in response to altered environmental and developmental contexts^11,12^. Many studies have focused on characterizing enhancers, the most studied type of CREs, involved in diseases, development, and cell- and tissue-type specificity, especially in mammalian models^11,13,14^. For example, in human renal cells the hyperosmotic induction of the sodium/myo-inositol cotransporter (***SMIT***) is mediated via several enhancers found in its 5’-untranslated region (UTR)^15^. In contrast to these findings in mammalian models, a comprehensive understanding of enhancer functions in fish exposed to salinity stress is still very limited. We have recently identified several copies of a CRE, the osmolality/salinity-responsive enhancer 1 (OSRE1) in the inositol monophosphatase (***IMPA1.1***) and *myo*-inositol phosphate synthase (***MIPS***) genes of *O. mossambicus*^16^. Enhancers such as OSRE1 are generally considered to function independent of whether they occur in the 5’ or 3’ regulatory regions or in introns^11,17^. Although most enhancers, including OSRE1 in *O. mossambicus* IMPA1.1 and MIPS genes^16^, are found in the 5’ regulatory region, intronic enhancers have been previously reported. For example, using human cell lines, Harris et al. have identified a tissue-specific enhancer in intron 1 of the cystic fibrosis transmembrane conductance regulator gene (***CFTR***)^18,19^. Another study has reported that enhancers located in intron 4 are responsible for differential expression of the Bone Morphogenetic Protein 6 gene (***Bmp6***), which underlies phenotypic differences between fresh water and seawater populations of threespine sticklebacks (*Gasterosteus aculeatus*)^20^.

The *glutamine synthetase* gene **(*GS)*** encodes an evolutionarily highly conserved enzyme that catalyzes the conversion of ammonia to glutamine. It is thought to be crucial for detoxification of ammonia as a part of nitrogen metabolism in diverse organisms including vertebrates^21–23^. Most studies on glutamine synthetase in fish, including euryhaline *O. mossambicus, O. niloticus* and *Oncorhynchus mykiss*, have focused on abundance or activity of glutamine synthetase in different organs such as intestine, muscle, liver and gills^24–28^. In addition to its function for nitrogenous waste detoxification in fish, glutamine synthetase also has an important function to maintain osmotic homeostasis. Glutamine synthetase produces glutamine, which can be accumulated in cells as a compatible organic osmolyte to offset the perturbing effects of hyperosmotic stress^6,29,30^. For example, in gills of the swamp eel (*Monopterus albus*) the induction of ***GS*** has been shown to promote accumulation of the compatible osmolyte glutamine during hyperosmotic stress^31^. However, little is known about transcriptional regulation of ***GS*** during salinity stress to adjust osmoregulation in euryhaline fish adapting to altered salinity.

Better knowledge of environmentally altered salinity effects on transcriptional regulation in fish is necessary to properly assess how global climate change that is predicted to accelerate salinization of many aquatic environments will impact the biodiversity and the future evolution of fish^32^. In addition, mechanistic insight into fish salinity (hyperosmotic) stress adaptation will contribute to improve aquaculture practices in brackish and increasingly saline environments^33^.

To contribute to better understanding adaptive mechanisms controlling fish osmoregulation, we investigated the transcriptional regulatory mechanism by which osmotic responsiveness is conferred to the *O. mossambicus* ***GS***. First, we analyzed whether the salinity-induced abundance increase of glutamine synthetase protein is based on transcriptional regulation. Then, an enhancer trapping reporter assay was used to identify the specific genomic regions that are responsible for transcriptional induction of ***GS*** during hyperosmolality.

## Results

### Hyperosmotic induction of glutamine synthetase is transcriptional and mediated by intron 1

Actinomycin D applied to OmB cells during exposure to hyperosmotic stress prevented glutamine synthetase production, which confirms that glutamine synthetase upregulation is mediated by transcriptional induction (Fig. 1 and Supplementary Fig. S1). Quantitation of glutamine synthetase abundance revealed a 4.65 ± 0.18 fold increase during hyperosmotic stress (mean ± s.e.m., p<0.0015, Fig. 1). This increase in glutamine synthetase abundance was completely abolished by including 10 mM actinomycin D in the media to yield a slight 0.85 ± 0.09 fold reduction during hyperosmotic stress (mean ± s.e.m, p=0.2726, Fig. 1).

**Figure 1.**
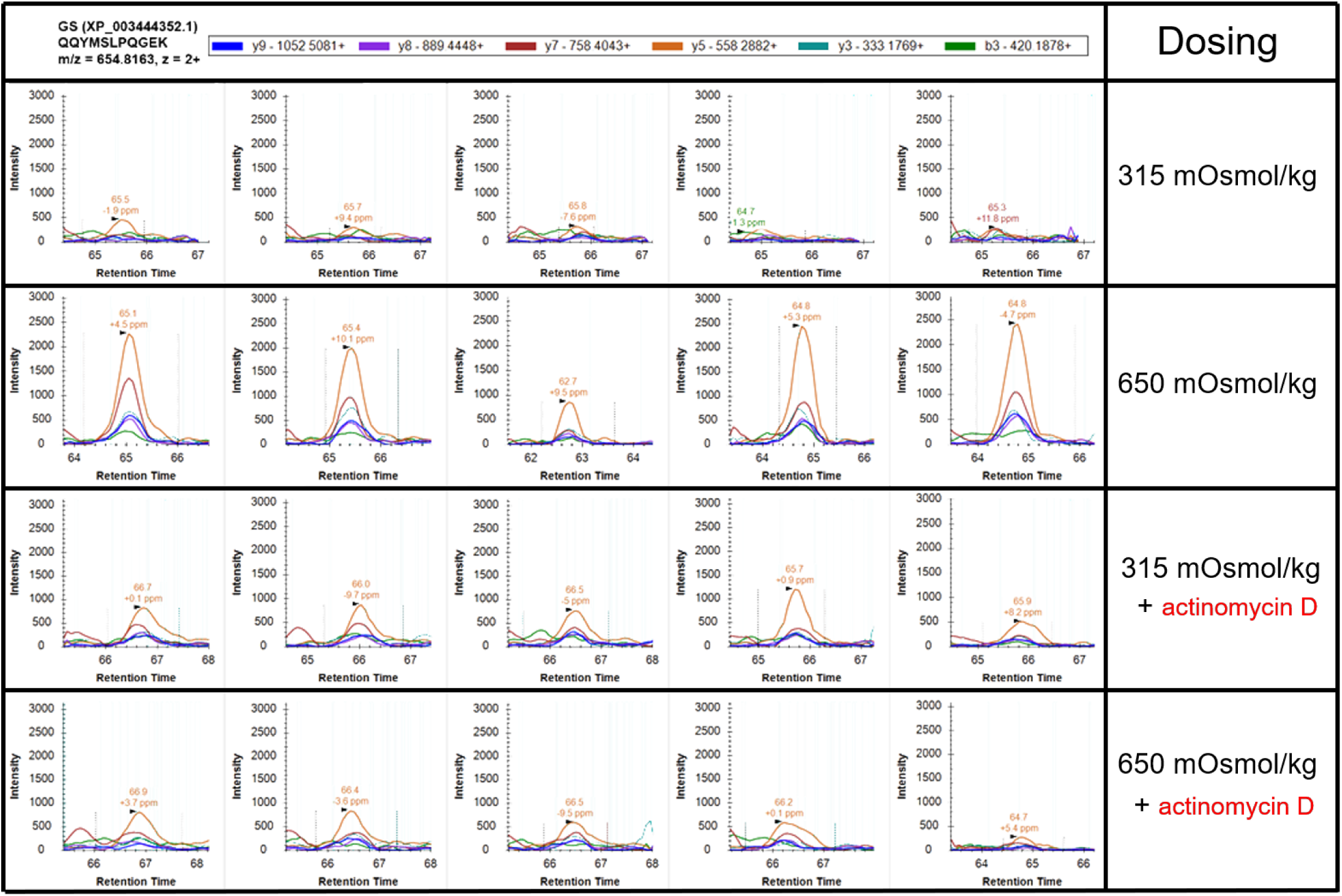
Targeted DIA-MS/Skyline protein quantitation of glutamine synthetase protein (GS, XP_003444352.1) in cells grown in four different medium conditions: isosmotic (315 mOsmol/kg), hyperosmotic (650 mOsmol/kg), isosmotic+ 10µM actinomycin D, hyperosmotic+10µM actinomycin D. Data for one of the four quantified peptides, QQYMSLPQGEK, is shown. Each treatment consisted of five biological replicates (from left to right).

The 3.4-kb 5’ regulatory sequence (RS), including the 5’-UTR, of the *O. mossambicus* ***GS*** was cloned, sequenced, and submitted to GenBank (GenBank Accession Number: MN631059). The start codon (SC) for translation was located in exon 2 (Fig. 2a). The first region tested for hyperosmotic enhancer activity was very long and spanned base pairs −2825 to the SC (+499). The corresponding plasmid construct with the first region inserted for a luciferase reporter assay is shown in Supplementary Fig. S2. This region conferred a 3.2±0.09(s.e.m)-fold (p<0.001) increase in luciferase reporter gene activity under hyperosmotic conditions relative to isosmotic controls (Fig. 2b). Iteratively narrowing this large region into successively shorter regions that had an identical 3’ end but differed at the 5’ end did not result in any loss of hyperosmotic induction of the reporter. These shortened constructs yielded 3.5± 0.18(s.e.m)-fold (p<0.001, −718 to SC), 3.4± 0.35(s.e.m)-fold (p<0.001, −257 to SC), 3.4± 0.24(s.e.m)-fold (p<0.001, −108 to SC), and 3.7±0.30(s.e.m)-fold (p<0.001, −60 to SC) reporter gene transcriptional induction, respectively (Fig. 2c). The shortest of these regions (559 bp) that is contained in all five constructs is composed of the core promoter, exon 1, and intron 1. These results suggest that the core promoter, exon 1, and/ or intron 1 are responsible for induction of the ***GS*** gene during hyperosmolality.

**Figure 2.**
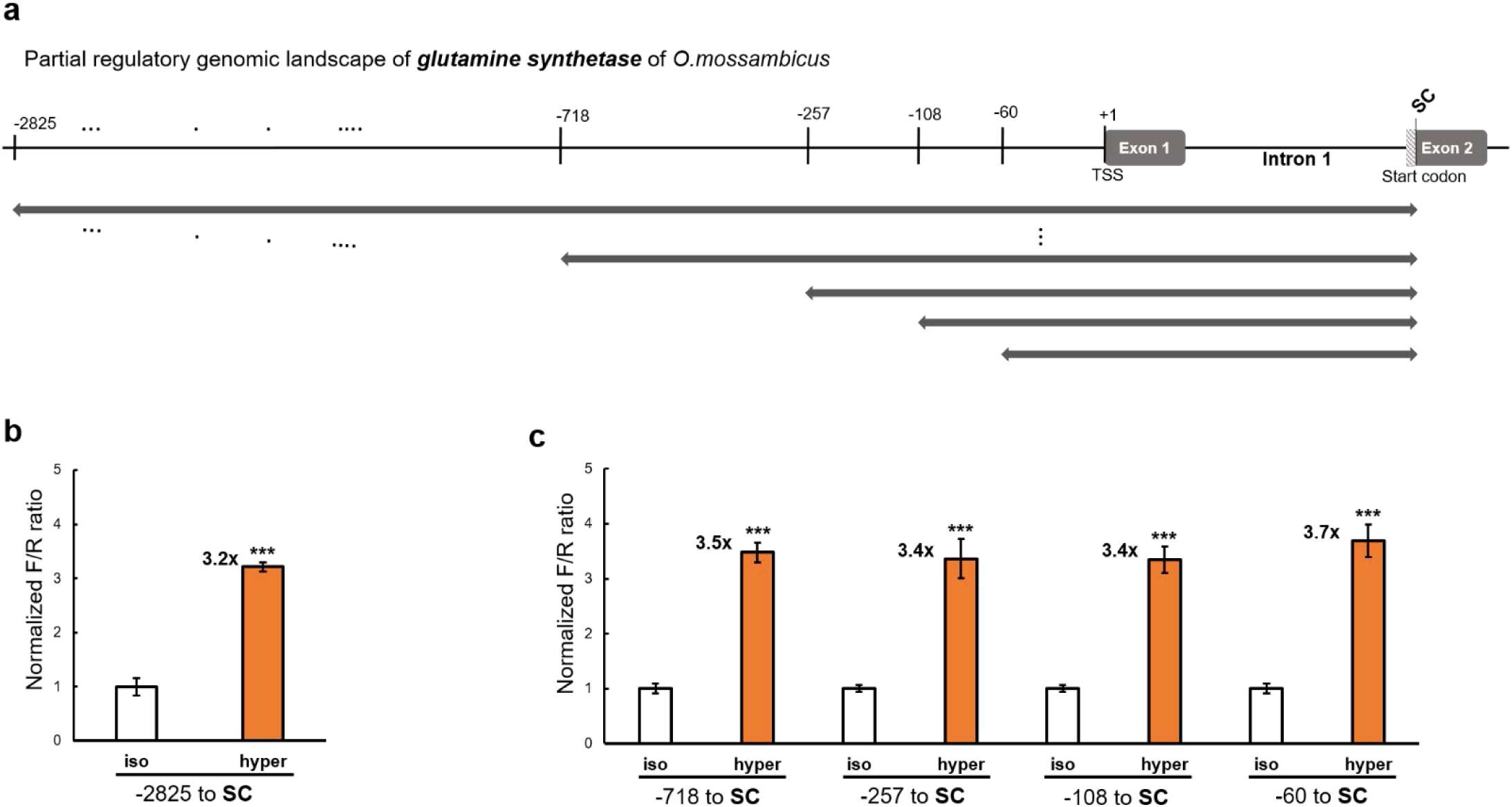
Narrowing of the osmotically regulated genomic region of the ***GS*** gene. **(a)** 3.4-kb long 5’-flanking genomic region and the 5’-UTR (including exon 1 and intron 1) of the *O. mossambicus* ***GS*** gene is illustrated. Numbers at the top indicate the genomic position relative to the transcription start site (TSS). The bars with arrows on both sides indicate the first set of genomic regions analyzed for hyperosmotic induction of reporter activity. The **SC** primer contains an NcoI restriction site at the translation start codon (SC); **(b and c)** Fold-change in luciferase reporter activity induced by hyperosmolality relative to isosmotic controls (**(b)** for the region −2825 to SC and **(c)** for successively shorter regions is shown. Normalized F/R ratio expresses inducible Firefly luciferase activity versus constitutive *Renilla* luciferase activity. This ratio was measured for both isosmotic (315 mOsmol/kg) and hyperosmotic (650 mOsmol/kg) conditions and normalized by setting isosmotic controls to one. The number of asterisks indicates the statistical significance of the hyperosmotic induction (p<0.001: ***, p<0.01: **, p<0.05: *, ns: not significantly different).

Because intron has recently been shown to confer transcriptional enhancement of several eukaryotic genes^34^, the role of intron 1 for the hyperosmotic ***GS*** induction was investigated further. First, intron 1 was excluded from all four shortened constructs to yield constructs that contain fragments whose 3’ end coincided with the end of exon 1 (+131 bp downstream of the transcription start site, TSS) (Fig. 3a). Removal of intron 1 completely abolished the hyperosmotic induction of reporter activity for all four of these constructs (−718 to +131, −257 to +131, −108 to +131, and −60 to +131) (Fig. 3b). This result demonstrates that intron 1 of ***GS*** is required for its hyperosmotic transcriptional induction. To test the hyperosmotic induction of intron 1 in a more physiological context using the endogenous rather than a heterologous core promoter we isolated the ***GS*** core promoter (GS-CP). For this purpose, a reporter plasmid containing the GS-CP (−257 to +131) was constructed (Fig. 3a). The functional GS-CP region (−257 to +131) was selected from four putative GS-CP regions because previous studies have shown that for many genes the functional promoter spans from approximately 250 bp upstream of the TSS to 100 bp downstream^35,36^. Deleting the region spanning −257 to −108 bp from the GS-CP abolishes GS-CP activity. In addition, we have identified three downstream promoter elements (DPEs) in the GS-CP by motif searching for the ‘RGWYVT’ consensus motif (Fig. 3a).

**Figure 3.**
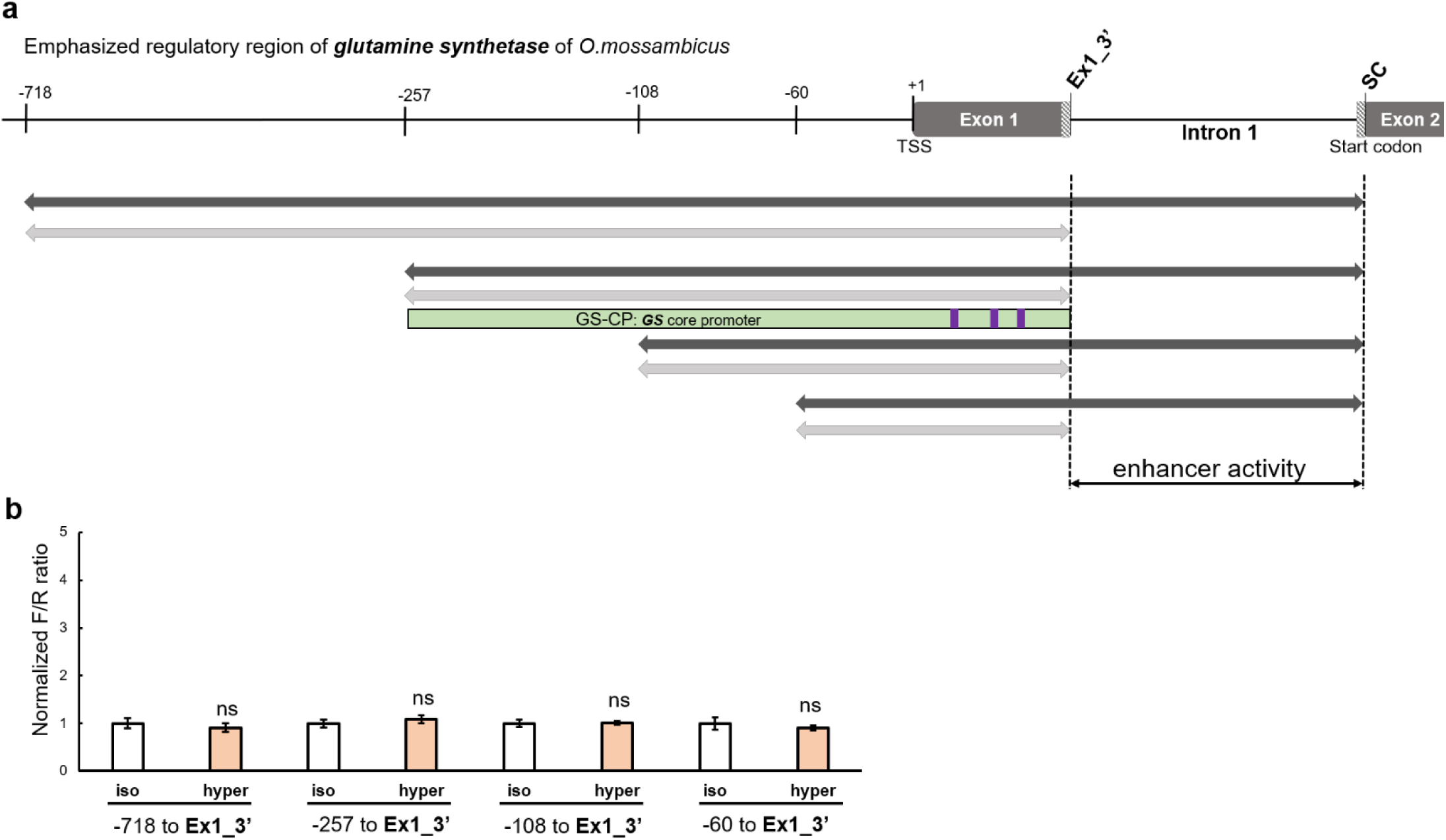
Identification of intron 1 as the genomic region necessary for hyperosmotic ***GS*** induction. **(a)** The genomic sequences used for reporter assays are shown. The **Ex1_3’** primer contains a NcoI restriction site at the 3’ end of exon 1. The grey bars indicate constructs that exclude intron 1 from the original constructs tested in Fig. 2(black bars). The light green bar presents the functional ***GS*** core promoter that is used for constructing the backbone reporter vector GS-CP+4.23. Three purple bars in the light green bar indicate downstream promoter elements (DPEs) that match to ‘RGWYVT’ motif. **(b)** Hyperosmotic induction of reporter activity is completely abolished when intron 1 is excluded from the luciferase constructs. The normalized F/R ratio expresses inducible Firefly luciferase activity versus constitutive *Renilla* luciferase activity. This ratio was measured for both isosmotic (315 mOsmol/kg) and hyperosmotic (650 mOsmol/kg) conditions and normalized by setting isosmotic controls to one. The number of asterisks indicates the statistical significance of the hyperosmotic induction (p<0.001: ***, p<0.01: **, p<0.05: *, ns: not significantly different). Figure layout and abbreviations are as outlined in Fig. 2.

### Intron 1 contains a single, functional copy of OSRE1

Systematic bioinformatics searches of the entire intron 1 sequence for the occurrence of a previously identified OSRE1 was performed by utilizing the OSRE1-consensus (DDKGGAAWWDWWYDNRB) and several specific OSRE1 sequences (incl. *O. mossambicus IMPA1.1*-OSRE1: AGTGGAAAAATACTAAG) that yielded high hyperosmotic induction of reporter activity in a previous study^16^. This approach enabled us to identify a single copy of OSRE1-like sequence (AGTGGAAAAATACAAC) in intron 1 of ***GS***. This ***GS***-OSRE1 was 16 bp long and almost identical (88%) to *IMPA1.1*-OSRE1, harboring only one gap and a single mismatch. ***GS***-OSRE1 was localized on the reverse strand (Fig. 4a).

**Figure 4.**
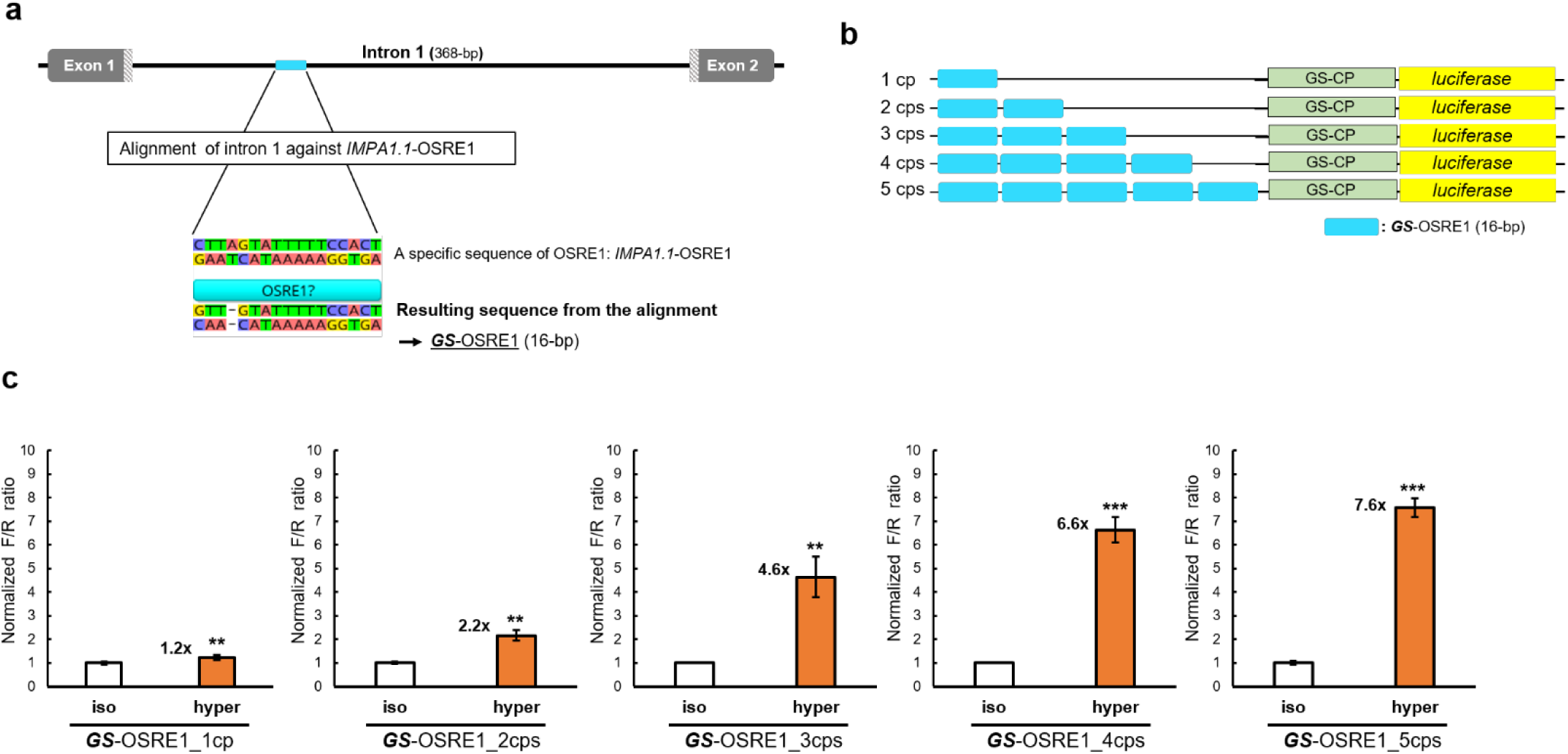
Identification of an osmoresponsive element (OSRE1) in intron 1 of ***GS*. (a)** Pairwise alignment of the ***GS*** intron 1 sequence against the 17 bp *IMPA1.1*-OSRE1 sequence yielded a match with 15 identical bases and 88.2 % of pairwise identity (one gap, one mismatch), which is referred to as ***GS***-OSRE1. **(b)** Reporter constructs containing different copy number of ***GS***-OSRE1 are depicted. **(c) *GS***-OSRE1 represents an inducible enhancer whose transcriptional potency is proportional to copy number. (cp = copy, cps = copies). The normalized F/R ratio expresses inducible Firefly luciferase activity versus constitutive *Renilla* luciferase activity. This ratio was measured for both isosmotic (315 mOsmol/kg) and hyperosmotic (650 mOsmol/kg) conditions and normalized by setting isosmotic controls to one. The number of asterisks indicates the statistical significance of the hyperosmotic induction (p<0.001: ***, p<0.01: **, p<0.05: *, ns: not significantly different).

To verify whether ***GS***-OSRE1 has functional activity as an enhancer element during salinity stress, a series of luciferase reporter plasmids driven by the endogenous GS-CP were constructed. Synthetic oligonucleotides harboring different numbers of copies of ***GS***-OSRE1 were used to validate its function as an osmoresponsive enhancer. Constructs containing either a single copy or up to five copies of ***GS***-OSRE1 were tested using dual luciferase reporter assays (Fig. 4b). Each of these constructs conferred hyperosmotic induction of reporter activity. Moreover, the extent of induction was proportional to the number of ***GS***-OSRE1 copies. However, a single copy yielded only a very small albeit significant degree of hyperosmotic induction 1.2±0.11(s.e.m)-fold (p<0.01). In contrast, two copies (2.2±0.23(s.e.m)-fold, p<0.01), three copies (4.6±0.86(s.e.m)-fold, p<0.01), four copies (6.6±0.55(s.e.m)-fold, p<0.001), and five copies (7.6±0.40(s.e.m)-fold, p<0.001) of ***GS***-OSRE1 yielded much greater hyperosmotic induction (Fig. 4c). These data demonstrate that ***GS***-OSRE1 functions as an osmoresponsive CRE during hyperosmotic stress. However, they also show that a single copy of ***GS***-OSRE1 is insufficient to explain the 3.4 to 3.7-fold hyperosmotic GS induction mediated by intron 1 (Fig. 2c).

After confirming the enhancer function of ***GS***-OSRE1 we refined the consensus sequence for OSRE1 by including the ***GS***-OSRE1 sequence in the consensus. This inclusion resulted in a change of the overall OSRE1 motif from DDKGGAAWWDWWY<u>D</u>NRB to DDKGGAAWWDWWY<u>N</u>NRB (Fig. 5).

**Figure 5.**
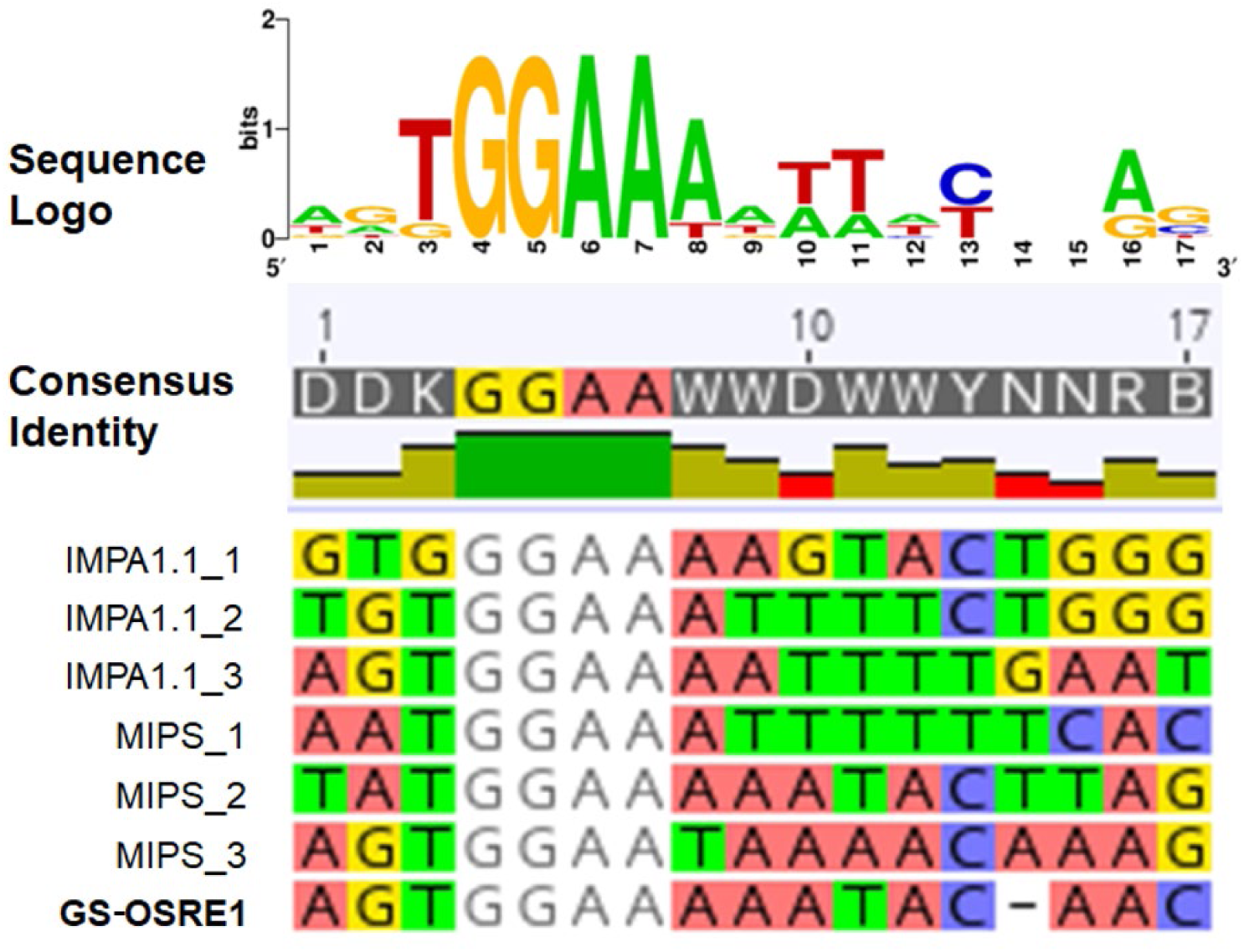
Refinement of osmolality/salinity-responsive enhancer 1 (OSRE1) consensus sequence. A multiple sequence alignment of ***GS***-OSRE1 with previously identified OSRE1 motifs in ***IMPA1.1*** and ***MIPS*** genes^16^ is shown. This alignment yields the refined consensus sequence DDKGGAAWWDWWYNNRB.

### Hyperosmotic induction of *GS* depends on the location of intron 1 and requires OSRE1

The dependence of hyperosmotic induction of ***GS*** on the location of intron 1 was investigated to address whether OSRE1-containing intron 1 behaves as a conventional position-independent enhancer^37^. Unexpectedly, when intron 1 was positioned downstream of the GS-CP (which represents its native genomic context) the hyperosmotic induction of reporter activity was much lower than when it was trans-positioned upstream of the GS-CP (3.4-fold vs. 9.9-fold, Fig. 6a and 6b). This result shows that intron 1-mediated transcriptional regulation of ***GS*** during salinity stress depends on the location of intron 1, which is atypical for conventional enhancers^17,37,38^. This atypical but pronounced position-dependency of intron 1 mediated enhancement represents a potential mechanism for evolutionary tuning of enhancer responsiveness via trans-positioning regulatory elements.

**Figure 6.**
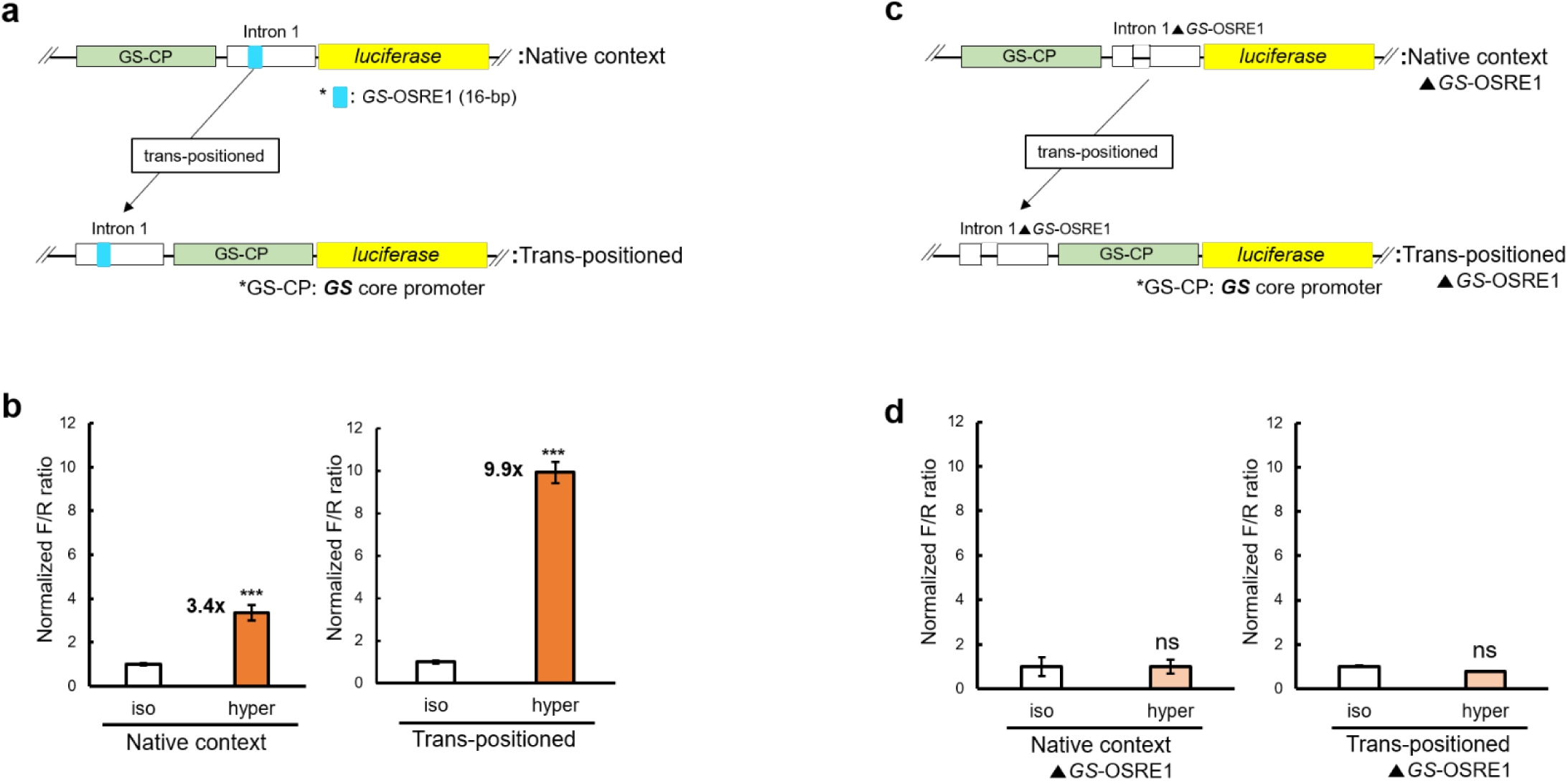
Two characteristics of the mechanism for hyperosmotic induction of ***GS*** by intron 1. **(a)** The genomic position of intron 1 was changed from downstream (native context) to upstream (trans-positioned) relative to the transcription start site (TSS). The light green bars indicate the ***GS*** core promoter (GS-CP). The light blue color indicates the ***GS***-OSRE1 element. **(b)** The corresponding reporter gene activity results are illustrated in **(a). (c, d)** The effect of selective deletion of ***GS***-OSRE1 from intron 1 (Intron 1▴***GS***-OSRE1) on reporter activity is shown. All reporter assays were carried out with reporter plasmid containing the GS-CP. Normalized F/R ratio expresses inducible Firefly luciferase activity versus constitutive *Renilla* luciferase activity. This ratio was measured for both isosmotic (315 mOsmol/kg) and hyperosmotic (650 mOsmol/kg) conditions and normalized by setting isosmotic controls to one. The number of asterisks indicates the statistical significance of the hyperosmotic induction (p<0.001: ***, p<0.01: **, p<0.05: *, ns: not significantly different).

In addition to establishing the position-dependency of intron 1 enhancement (Fig. 6a and 6b) and functionally validating OSRE1 (Fig. 4), we investigated whether ***GS***-OSRE1 is necessary for the enhancer function of intron 1. To test whether the presence of ***GS***-OSRE1 is essential for intron 1-mediated hyperosmotic transcriptional induction of ***GS*** the 16 bp OSRE1 sequence was deleted from intron 1. The rationale for this experiment was that, although a single copy of ***GS***-OSRE1 was insufficient to account for the hyperosmotic induction of the ***GS*** gene (Fig. 4c), it may still be required as an essential component triggering the formation an inducible transcription factor complex. Two luciferase reporter plasmids with deletions of ***GS***-OSRE1 were constructed and tested for luciferase activity in OmB cells exposed to iso- and hyperosmotic media (Supplementary Fig. S3). One of these constructs harbored intron 1 downstream of the TSS in its native context and the other contained intron 1 trans-positioned upstream of the TSS (Fig. 6c). Selectively deleting ***GS***-OSRE1 (16 bp) from intron 1 completely abolished the hyperosmotic transcriptional induction conferred by intron 1, independent of the location of intron 1 (Fig. 6d). This result demonstrates that ***GS***-OSRE1 is necessary for the enhancer activity of ***GS*** intron 1.

## Discussion

In this study, we discovered a novel molecular mechanism where intron 1 harboring an osmoresponsive CRE (***GS***-OSRE1) positively regulates transcription of the tilapia ***GS*** under hyperosmotic stress. Furthermore, we identified that intron 1-mediated hyperosmotic ***GS*** induction requires the OSRE1 element and that its enhancer activity depends on the location of intron 1. The molecular mechanism of intron 1 enhancement of ***GS*** transcription during hyperosmotic stress differs from that of conventional context-inducible transcriptional enhancers, many of which have been previously shown to function independent of orientation or distance (relative position) to the TSS^17,39^.

Previously, we analyzed the 5’ RS of *O. mossambicus* ***IMPA1.1*** and ***MIPS*** genes and identified six osmotically responsive CREs with a common 17 bp consensus motif, which we named OSRE1^16^. All of these OSRE1 elements were located between −232 and +56 bp relative to the TSS in both genes. The first osmotically responsive enhancers were identified in mammalian cell lines and named Tonicity responsive element (TonE)^40^ and, alternatively, osmotic response element (ORE)^41^. Bai et al. have also characterized a distinct osmotic-responsive element (OsmoE) in a mouse kidney cell line and revealed its genomic locus to be further upstream (−808 to −791 bp relative to the TSS) in the *NHE-2* gene encoding the Na^+^/H^+^ exchanger-2 ^42^. In addition to the discovery of OsmoE, this study also identified a TonE-like element far upstream (−1201 to −1189 bp) in the same gene^42^. Therefore, our initial attempt to identify an osmotically responsive CRE in the ***GS*** utilized the 3.4-kb region of 5’ RS spanning from –2825 to +526 bp. Our tilapia study and these previous observations in mammalian model systems suggest (with rare exceptions) that osmoresponsive CREs are located preferentially very close (within a few hundred bp) to the TSS. This knowledge informs comparative studies and future searches for osmoresponsive CREs in other genes and/ or species.

We elucidated that intron 1 in combination with the endogenous GS-CP mediates transcriptional induction of the ***GS*** under salinity stress. Introns have been shown to boost gene expression in numerous ways including by providing binding sites for transcription factors, regulating the rate of transcription, promoting nuclear export, and stabilizing transcripts^34^. Several studies of plant species have identified a positive effect of introns on transcription or mRNA accumulation in a constitutive rather than context-dependent manner^43,44^. Moreover, most reported cases of intron-mediated transcriptional enhancement are stimulus-independent^45,46^. Only a small number of studies has thus far investigated the stimulus-responsiveness of introns. However, some previous studies using different cell lines have shown that stimulus-dependent transcriptional regulation of a variety of genes is mediated by intron 1^47–49^. For example, in a human breast cancer cell model, intron 1 of the *ERBB2* proto-oncogene (***ERBB2***) contains a 409 bp sequence that mediates ***ERBB2*** transcriptional changes in response to oestrogens^47^. These previous studies reporting stimulus-dependent intron 1 mediated enhancement are consistent with our finding that intron 1 enhances ***GS*** transcription during hyperosmotic stress. Therefore, our study provides evidence that introns, which have often been regarded as “junk DNA” that is spliced out during mRNA processing, represent functional genomic targets for evolutionary adaptation to environmental changes.

Our observation that the degree of hyperosmotic transcriptional ***GS*** induction mediated by intron 1 is position-dependent suggests that the corresponding mechanism is distinct from typical CRE-mediated enhancement. A position-dependent effect of an intronic enhancer was also reported for intron 2 of the human beta-globin gene demonstrating that changes in the location of intron 2 relative to the promoter alters transcriptional activity three-fold^38^. This result is very similar to the three-fold change in transcriptional activity observed in our study when the location of ***GS*** intron 1 was altered. Moreover, the position dependence of intron-mediated enhancement (IME) has been well documented in plants^50^.

In other cases reported for IME, however, intron 1 has been shown to act independent of its location^48,49^. It is likely that sequence rearrangements around the TSS result in conformational changes of the transcriptional machinery, which then affects transactivation efficiency^51^. Therefore, it is possible that intron 1 trans-positioning changes the structural conformation of the transcriptional machinery in a way that increases transactivation. The location of intron 1 in a position that does not maximally enhance transactivation suggests that evolution has favored moderate over strong transcriptional induction of ***GS*** during hyperosmolality. Otherwise, transposition of the CRE elements included in intron 1 upstream of the TSS would have been evolutionarily favored. Possible reasons for limiting the extent to which ***GS*** is induction during hyperosmotic stress are as follows: Glutamine synthetase abundance during hyperosmotic stress may represent a compromise between its ability to produce a compatible organic osmolyte (glutamine) on the one hand and its consumption of energy (ATP) on the other hand. In most organisms including fish, glutamine synthetase is an essential enzyme that mediates bidirectional biochemical reactions, ammonia assimilation and glutamine biosynthesis^52,53^. Thus, a moderate increase of glutamine synthetase abundance during hyperosmotic stress may be evolutionarily favored as the most cost-effective strategy during salinity stress.

This study demonstrates that the ***GS***-OSRE1 element in intron 1 is essential for transcriptional induction during hyperosmotic stress. We interpret these results as evidence that the OSRE1 element serves as a critical binding site for a hyperosmolality-inducible transcription factor. Furthermore, our results suggest that a combination of inducible transcription factors is necessary for promoting transcriptional enhancement since a single copy of ***GS***-OSRE1 outside its native intron 1 sequence context was inefficient for enhancing transactivation. We conclude that other, yet to be identified CREs, are present in intron 1 that interact with OSRE1 to result in transcriptional enhancement^54^. Such combinatorial interactions between different CREs and corresponding transcription factors are common^55,56^. One important focus of future research will be to characterize such complexes and their interactions.

In conclusion, ***GS*** intron 1 was revealed to contain a single OSRE1 (***GS***-OSRE1) and to enhance transcriptional induction of ***GS*** in a tilapia (*O. mossambicus*) cell line exposed to hyperosmolality. The mechanism for this transcriptional enhancement of ***GS*** expression during hyperosmolality has two characteristics: 1. Its extent is dependent on the location of intron 1 relative to the TSS, 2. It requires ***GS***-OSRE1 for intron 1 enhancer function. Furthermore, our data strongly suggest that the previously identified osmoresponsive CRE OSRE1 consensus sequence can be used for bioinformatics screening approaches that identify candidate OSRE1 sequences on a genome-wide bases^57^. Identification of the transcription factor(s) that bind to ***GS***-OSRE1 and potential other osmoresponsive CREs in intron 1 represents an intriguing future task to understand the process by which osmotic stress signals are perceived and transduced to regulate the expression of genes that compensate for salinity stress in euryhaline fish.

## Methods

### Cell culture

The tilapia OmB cell line^58^ was used for all experiments and luciferase reporter assays. OmB cells were maintained in L-15 medium containing 10 % (vol/vol) fetal bovine serum (FBS) and 1 % (vol/vol) penicillin-streptomycin at 26 °C and 2 % CO_2_. Using a large supply of OmB cell superstock ^16^ (passage 15; P15), all experiments were carried out on OmB cells between P18 to P26. Cells were passaged every 3-4 d using a 1:5 splitting ratio. For applying hyperosmotic stress to OmB cells, hyperosmotic (650 mOsmol/kg) medium was prepared using hypersaline stock solution (osmolality: 2820 mOsmol/kg). This stock solution was made by adding an appropriate amount of NaCl to regular isosmotic (315 mOsmol/kg) L-15 medium. The hypersaline stock solution was then diluted with isosmotic medium to obtain hyperosmotic medium of 650 mOsmol/kg. Medium osmolality was always confirmed using a freezing point micro-osmometer (Advanced Instruments).

### Proteomics

Sample preparation by tryptic in solution digestion, data-independent acquisition (DIA) and targeted proteomics were performed as previously described using a nanoAcquity UPLC (Waters), an ImpactHD mass spectrometer (Bruker), and Skyline^59^ targeted proteomics software^16^. Three peptides of GS that are identical in sequence in *O. mossambicus* and *O. niloticus* (NCB Accession # XP_003444352.1) were used for quantitation (Supplementary Fig. S1). Three proteins, represented by at least three peptides each, were used for normalization (fatty acid-binding protein, NCB Accession # XP_003444095.3, beta-tubulin, NCB Accession # XP_003455078.1, and actin 2, NCB Accession # XP_003455997.3).

### Cloning

Total genomic DNA was extracted from spleen tissue of Mozambique tilapia (*O. mossambicus*) using the PureLink Genomic DNA mini Kit (Invitrogen). PCR primers were designed with Geneious 11.0.3 (Biomatters) using the *O. niloticus* glutamine synthetase (NCB Accession # XM_003444304.4 and XP_003444352.1) genomic sequence as a template. A CCCCC spacer followed by a restriction enzyme recognition site was added to the 5’ end of each primer. The restriction enzymes KpnI, SacI, HindIII, and NcoI (New England BioLabs) were used to clone PCR amplicons representing genomic regions of the ***GS*** gene into pGL4.23 vector. Platinum PCR Supermix (Thermo Fisher Scientific) and/or Q5^®^ High-Fidelity DNA Polymerase (New England BioLabs) were used to amplify DNA fragments longer than 2 kb. For fragments < 2 kb, PCR Master Mix 2x (Promega) was used. PCR was conducted as follows: initial denaturation at 94°C for 3 min followed by 35 cycles of 94°C for 30 s, annealing: 48-60° for 30 s, elongation: 72°C for 0.5-2 min, and 72°C for 15 min. Annealing temperature and extension time were set according to the chemical features of the primers and the lengths of amplicons. PCR products were confirmed by agarose gel electrophoresis and sequentially either purified using the PureLink PCR Purification Kit (Thermo Fisher Scientific) or gel-extracted using the QIAquick^®^ Gel Extraction Kit (Qiagen). Specific primers were designed for the translation start site (start codon, **SC**, +499) and the 3’ end of exon 1 (**Ex1_3’**, +131). The **SC** and **Ex1_3’** primers included a NcoI restriction site that was already present in the wildtype ***GS*** donor sequence and in the pGL4.23 acceptor reporter plasmid. Therefore, genomic regions of interest that terminate at the SC and Ex1_3’ sites could be cloned without changing any wildtype sequence. All amplified ***GS*** gene fragments were double-digested with two enzymes (combinations of KpnI, SacI, HindIII, and NcoI). Restriction enzyme digestion was conducted in 10 μL reaction buffer (CutSmart®Buffer and NEBuffer™ 1.1) containing 2 μL (10 U/μL) of each restriction enzyme, 0.5-2 μg of purified PCR product, and nuclease-free H_2_O ad 100 μL. After overnight incubation at 37°C, reactions were stopped by 20 min incubation at 80°C. Digested inserts and vectors were purified using the PureLink™ Quick PCR Purification Kit (Thermo Fisher Scientific) and ligated to produce recombinant constructs using T4 DNA ligase (Thermo Fisher Scientific). Ligation reactions contained 50 ng of vector, 10-20 ng of insert (depending on its size to yield a 1:3 or 1:5 molar ratio), 2 μL of ligase buffer, 1 μL of T4 ligase (1 U/μL) and nuclease-free H_2_O ad 20 µL. Ligation proceeded at 25°C for 6h. The ligation products were transformed into 10-beta-competent *E. coli* (New England Biolabs) as follows: First, a 50 μL aliquot of bacteria was thawed on ice for 5 min, then 10 μL of bacterial suspension was added to 1.5 μL of a single ligation reaction. Second, the mixture was kept on ice for 30 min, exposed to heat shock (42°C) for exactly 30 s, and placed back on ice for 5 min. Third, 190 μL of super optimal broth with catabolite repression medium (SOC, Thermo Fisher Scientific) was added and transformed bacteria were incubated at 250 rpm and 37°C for 60 min. After transformation, 30 μL of the bacterial solution was spread onto a pre-warmed (37°C) LB-ampicillin plate, which was used for single colony picking and colony PCR on the next day to confirm the presence of intended inserts. For colony PCR, tubes containing a bacterial clone were first quick-vortexed, then heated at 95°C for 15 min and quick-spun to remove debris. Three μL of the supernatant were mixed with forward and reverse primers that flank the corresponding insert. Colony PCR conditions were the same as described above and amplicons were checked by agarose gel electrophoresis. Colonies that contained an insert of the expected size were chosen for plasmid purification. Each bacterial colony was inoculated into liquid LB medium and grown for 16-18 h to maximize plasmid yield. Liquid cultures were harvested and purified according to manufacturer’s protocol using endotoxin-free PureLink™ Quick Plasmid Miniprep Kit (Thermo Fisher Scientific). Insert sequences in purified DNA constructs were verified by Sanger sequencing at the University of California, Davis DNA Sequencing Facility before using the corresponding constructs for transient transfection into tilapia OmB cells.

### Enhancer trap reporter assays

Enhancer trapping assays were performed according to the protocol previously reported by our laboratory^16^. To produce a backbone luciferase vector harboring the endogenous functional promoter of the ***GS***, the functional ***GS*** core promoter region (GS-CP, −257 to +131, Fig. 3a) was cloned into upstream of the firefly (*Photinus pyrails*) luciferase gene in pGL4.23 vector (GenBank Accession Number DQ904455.1, Promega) and verified that it has constitutive activity but is not hyperosmotically inducible. The resulting reporter plasmid was named GS-CP+4.23. The GS-CP region was amplified using a forward primer that included a HindIII restriction site and a reverse primer that included a NcoI restriction site. The GS-CP region and pGL4.23 plasmid were digested with the same pair of restriction enzymes and followed by ligation. Cloning, purification, and sequence-validation were conducted as described in the cloning procedure.

The GS-CP+4.23 plasmid was used in combination with *hRluc (Renilla reniformis*) luciferase control plasmid pGL4.73 (GenBank Accession Number AY738229.1, Promega). Co-transfection of tilapia OmB cells with this control plasmid was used to normalize for variability of transfection efficiency and cell number. One day prior to co-transfection OmB cells were seeded in 96-well plates (Thermo Fisher Scientific) at a density of 2 x 10^4^ cells per well. Co-transfection was performed when cells reached 80% to 90% confluency. Co-transfection was performed with ViaFect (Promega) reagent using previously optimized conditions ^16^. Cells were allowed to recover for 24h after transfection before being dosed in either isosmotic (315 mOsmol/kg) or hyperosmotic (650 mOsmol/kg) media for 72h. Dual luciferase activity was measured in 96-well plates using a GloMax Navigator microplate luminometer (Promega). Four biological replicates were used for each experimental condition. All luciferase raw measurements were adjusted for transfection efficiency by normalizing the firefly luciferase activity to *Renilla* luciferase activity. They were expressed as fold-change in hyperosmotic media relative to isosmotic controls. One-way ANOVA was performed to assess statistical significance of the data and calculate p values using R package software (http://www.R-project.org/).

### Bioinformatics sequence analysis

Intron 1 was searched for the occurrence of an OSRE1 consensus motif using a bioinformatics approach. For this purpose, Geneious 11.0.3 (Biomatters) was used. Both strands, sense and antisense, were searched. Sequence similarity searches were conducted by using the overall OSRE1-consensus sequence (DDKGGAAWWDWWYDNRB) as well as several experimentally validated and previously identified variants of OSRE1 sequences, including the 17 bp sequence AGTGGAAAAATACTAAG (*IMPA1.1*-OSRE1), as templates^16^.

### Synthetic oligonucleotide annealing and GeneStrands synthesis

The effect of ***GS***-OSRE1 copy number variation and ***GS***-OSRE1 deletion on hyperosmotic reporter activity was analyzed. Synthetic oligonucleotides containing different copy numbers of ***GS***-OSRE1 were produced by oligonucleotide annealing (Eurofins Genomics). ***GS***-OSRE1 constructs containing one, two, three, four and five copies were generated. Forward and reverse PCR primers for amplifying each synthetic oligonucleotide were designed to contain SacI and HindIII restriction sites to enable subsequent cloning into GS-CP+4.23 vector (Supplementary Table S1). Synthetic oligonucleotides harboring more than three copies of ***GS***-OSRE1 or mutated intron 1 (Intron 1▲***GS***-OSRE1) were longer than 100 bp. These longer inserts were synthesized using the GeneStrands method (Eurofins Genomics). Subsequently, each insert was separately cloned into GS-CP+4.23 luciferase reporter vector. After cloning into the reporter plasmid, the proper sequences of all inserts were verified by Sanger sequencing. These constructs were used to assess the effect of ***GS***-OSRE1 copy number and deletion of ***GS***-OSRE1 from intron 1 on reporter activity under hyperosmotic (650 mOsmol/kg) conditions relative to isosmotic controls (315 mOsmol/kg).

## Data Availability

All data generated or analyzed during this study are included in this manuscript and its Supplementary information files. Sequence data for the 5’ RS of the *O. mossambicus* ***GS*** gene investigated in this study can be found in GenBank with Accession Number: MN631059. The DIA assay library, results, and metadata for glutamine synthetase quantitation are publicly accessible in the targeted proteomics database Panorama Public^60^ at the following link: https://panoramaweb.org/lUknq6.url.

## Acknowledgements

This investigation was supported by National Science Foundation (NSF) grant IOS-1656371.

## Author contributions statement

D.K. conceived the project, C.K. performed the experiments, conducted the bioinformatics analyses, and analyzed the results and C.K. and D.K. wrote the manuscript.

## Additional information

The authors declare no competing interests.

