## Supplementary Materials for "An osmolality/ salinity-responsive enhancer 1 (OSRE1) in intron 1 promotes salinity induction of tilapia glutamine synthetase"

**Supplementary Figure S1**. Additional targeted SWATH-MS/Skyline protein quantitation data of GS protein in cells grown in four different medium conditions (Dosing: isosmotic (315 mOsmol/kg), hyperosmotic (650 mOsmol/kg), isosmotic+ 10µM actinomycin D, hyperosmotic+10µM actinomycin D). Two different peptides of glutamine synthetase (EEGEEPANYSK, RPSANCDPYAVTEALVR) are shown with five biological replicates for each dosing condition.

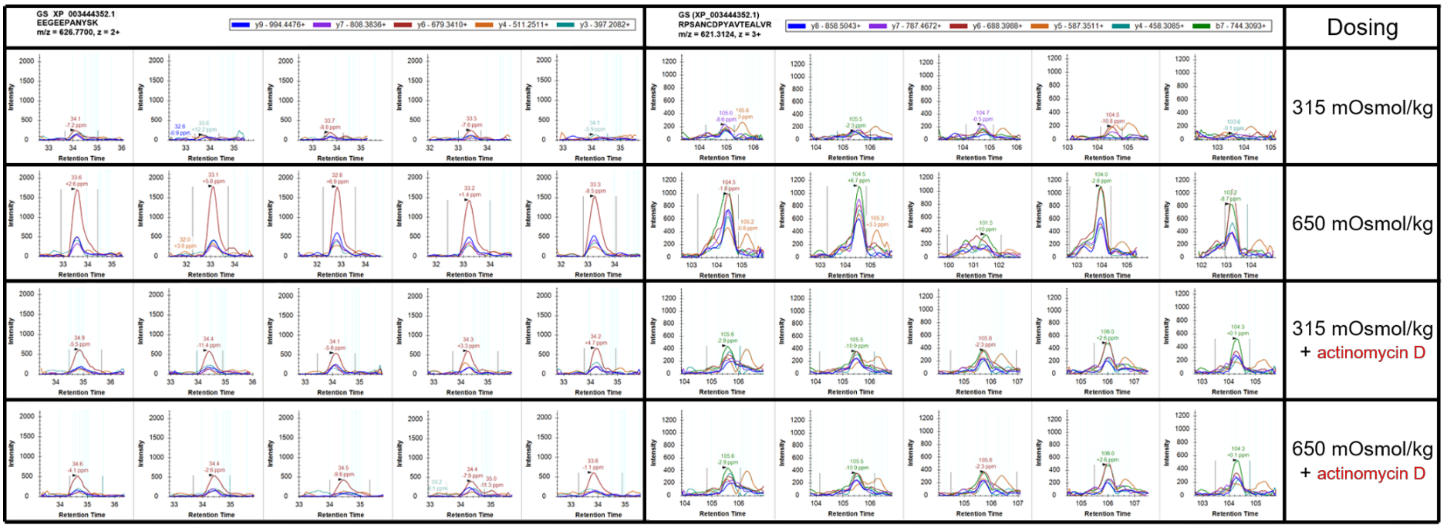


**Supplementary Figure S2**. Plasmid map of the initial luciferase reporter construct consisting of pGL4.23 vector (blue) and an insert representing the 3.4-kb 5' regulatory sequence (RS) of ***GS*** (orange, -2825 to +499). The 3' end of the insert (+499) represents the start codon (SC). Exon 1, the start codon at the beginning of Exon 2 (dark grey), intron 1 (green) of ***GS*** and the luciferase reporter gene (yellow) are indicated.

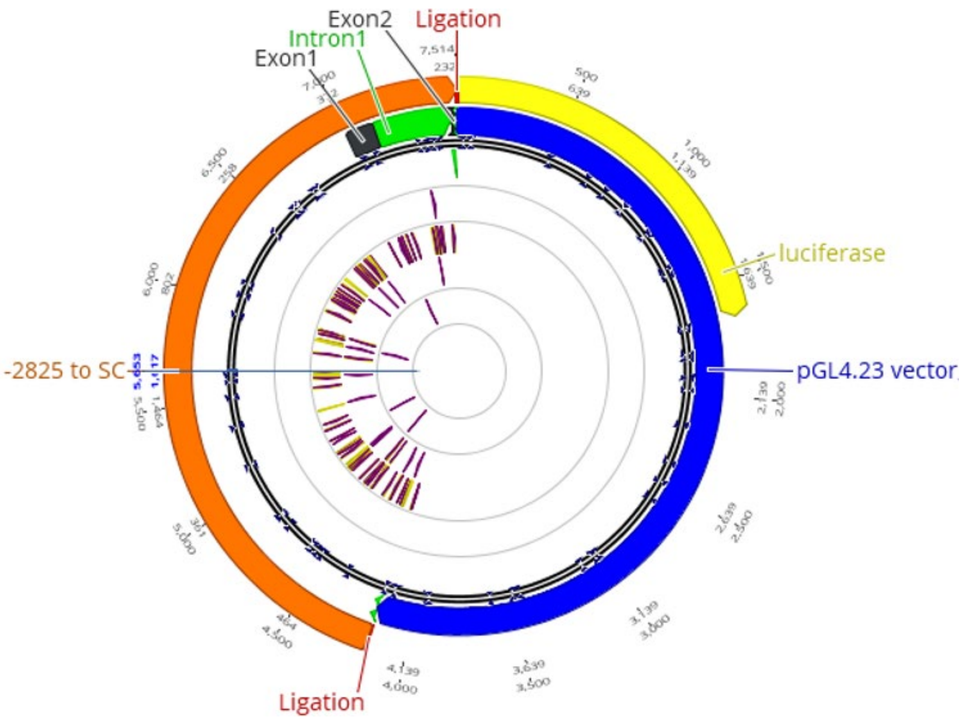

**Supplementary Figure S3**. Sequencing result confirming selective deletion of ***GS***-OSRE1 from intron 1 in the reporter contsruct.

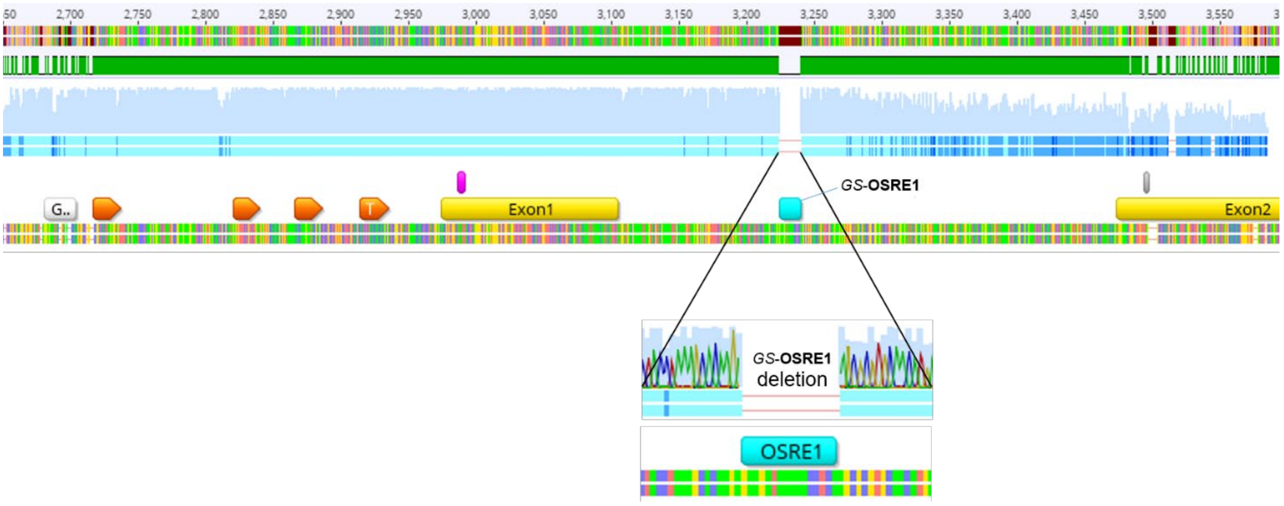


**Supplementary Table S1**. Synthetic oligonucleotide sequences containing different copy numbers of ***GS***-OSRE1 (SacI and HindIII restriction sites are also included) and the corresponding Forward & Reverse primer sequences for their amplification. Red and black color variation separates multiple ***GS***-OSRE1 copies.

*GS*-OSRE1: GTTGTATTTTTCCACT

| Number of copy | Whole oligonucleotide sequence (bp length) | Forward primer Sequence | Reverse primer sequence |
|---|---|---|---|
| 1 | CCCCCGAGCTCCATGCATGCATGCTGTGTTGTATTTTTCCACTTGCATGCATGCATGCCGAGCTAAGCTTGGGGG<br>(75 bp) | CCCCCGAGCTCCATGCATGCATGCTGTGTTGTATTTTTCCACTTGCATGCATG | CCCCCAAGCTTAGCTCGGCATGCATGCATGCAAGTGGAA |
| 2 | CCCCCGAGCTCCGTTGTATTTTTCCACTACTGGTTGTATTTTTCCACTCGAGCTAAGCTTGGGGG<br>(65 bp) | CCCCCGAGCTCCGTTGTATTTTTCCACTACTGGTTGTATTTTTCC | CCCCCAAGCTTAGCTCGAGTGGAAAAATACAACCAGTAGTGG |
| 3 | CCCCCGAGCTCTGTGTTGTATTTTTCCACTTGCACTGTGTGTTGTATTTTTCCACTTGCACTGTGTGTTGTATTTTTCCACTTGCAAGCTTGGGGG<br>(96 bp) | CCCCCGAGCTCTGTGTTGTATTTTTCCACTTGCACTGTGTGTTGTATTTTTCCAC | CCCCCAAGCTTGCAAGTGGAAAAATACAACACACAGTGCAAGTGGAAAAATACAACACACA |
| 4 | CCCCCGAGCTCTGTGTTGTATTTTTCCACTTGCACTGTGTGTTGTATTTTTCCACTTGCCGAGCTTGTGTTGTATTTTTCCACTTGCACTGTGTGTTGTATTTTTCCACTTGCAAGCTTGGGGG<br>(124 bp) | GeneStrands synthesis (Eurofins Genomics) | |
| 5 | CCCCCGAGCTCTGTGTTGTATTTTTCCACTTGCACTGTGTGTTGTATTTTTCCACTTGCACTGTGTGTTGTATTTTTCCACTTGCACTGTGTGTTGTATTTTTCCACTTGCACTGTGTGTTGTATTTTTCCACTTGCAAGCTTGGGGG<br>(148 bp) | GeneStrands synthesis (Eurofins Genomics) | |
